# Telomere protein arrays stall DNA loop extrusion by condensin

**DOI:** 10.1101/2023.10.29.564563

**Authors:** Brian T. Analikwu, Alice Deshayes, Jaco van der Torre, Thomas Guérin, Allard J. Katan, Claire Béneut, Roman Barth, Jamie Phipps, Vittore Scolari, Xavier Veaute, Christopher Barrington, Didier Busso, Frank Uhlmann, Karine Dubrana, Stefano Mattarocci, Cees Dekker, Stéphane Marcand

## Abstract

DNA loop extrusion by SMC proteins is a key process underlying chromosomal organization. It is unknown how loop extruders interact with telomeres where chromosome ends are covered with a dense array of tens of neighboring DNA-binding proteins. Using complementary *in vivo* and *in vitro* single-molecule approaches, we study the interaction between loop-extruding condensin and arrays of Rap1, the double-stranded-DNA-binding telomeric protein of *Saccharomyces cerevisiae*. We show that dense linear Rap1 arrays can completely halt DNA loop extrusion, where the blocking efficiency depends on the array length and the DNA gap size between neighboring proteins. In cells, Rap1 arrays in the chromosome are found to act as contact insulators and to accumulate condensin at their borders, with direct implications for the resolution of dicentric chromosomes produced by telomere fusions. Our findings show that linear arrays of DNA-bound proteins can efficiently halt DNA loop extrusion by SMC proteins, which may impact a wide range of cellular processes from telomere functions to transcription and DNA repair.

## MAIN

Telomeres are essential protein-DNA complexes that ensure that chromosome ends escape the pathways acting on broken DNA ends. They consist of long stretches of DNA with repetitions of short motifs tightly covered by sequence-specific DNA-binding proteins such as Rap1 in budding yeast^1–4^. Because of the tight packing, access to telomere DNA is restricted for DNA-processing events such as transcription, DNA repair, and replication^5–12^.

Here, we aim to shed light on the handling of such a tight DNA coverage at telomeres by a key organizer of chromosomal structure, the SMC complex (Structural Maintenance of Chromosomes) condensin. SMC complexes are motor proteins that extrude loops of DNA to organize chromatin into higher-order structures^13–20^. Condensin compacts chromosomes during mitosis via DNA loop extrusion^21–25^ and is essential to chromosome segregation^26,27^. Condensin consists of two ATPase SMC coiled-coil subunits (Smc2 and Smc4), a kleisin (Brn1 in budding yeast), and two HEAT-repeat subunits (Ycs4 and Ycg1 in budding yeast). Yeast condensin acts as a monomeric protein complex that anchors DNA at the Brn1-Ycg1 interface and extrudes DNA into a loop from this anchoring point^15,18,28,29^. DNA loop extrusion is driven by ATP-dependent conformational changes and multiple dynamic DNA-protein contacts (refs. 18,30,31, and Dekker et al, Science, to appear on Nov.10).

It is currently intensely studied whether loop extrusion by condensin and other SMC complexes can be blocked by DNA-binding proteins that may act as roadblocks for loop extrusion^32–36^. The DNA-binding protein CTCF, known to demarcate the boundaries of topologically associated domains (TADs)^34,37^, was recently shown to block the SMC complex cohesin in a direction- and force-dependent manner through specific chemical interactions^33^. In the absence of a biochemical interaction, SMC complexes were, by contrast, found to be remarkably efficient at passing isolated physical roadblocks on the DNA *in vitro*^36^. However, in cells, chromosome-bound roadblocks are often not present as single obstacles at low density. For instance, RNA polymerases have been reported to stall SMC complexes at highly transcribed genes, perhaps as a consequence of DNA coverage by so-called polymerase trains^32,38–41^.

Our previous work suggested that condensin may stall at telomeres^12^. Upon studying dicentric chromosome breakage in *Saccharomyces cerevisiae*, we found that dicentrics resulting from accidental telomere-telomere fusions preferentially broke at the fusion points^42^ during abscission (septum closure in yeast)^12,43,44^. This restored the parental karyotype, therefore providing a backup pathway for telomere protection and genome stability. Breakage at telomere fusions requires two specific actors, namely condensin and the telomere DNA-binding protein Rap1. Condensin stalling by arrays of Rap1 might favor their capture at the abscission point, which would explain dicentric preferential breakage at telomere-telomere fusions^12^.

Here we employ both *in vitro* single-molecule and *in vivo* approaches to directly address how arrays of Rap1 impact condensin-driven loop extrusion. Dense and tightly bound telomeric repeats provide a unique setting to systematically and mechanistically study this interaction. We show that telomere Rap1 arrays inserted exogenously within a chromosome lead to an accumulation of condensin at their borders yielding a local boundary to chromatin compaction. By studying encounters between individual loop-extruding condensin complexes and Rap1 arrays in single-molecule visualizations, we show how ∼100 nm arrays can stall condensin by physically blocking the loop extrusion with near-100% efficiency. Stalling is modulated by DNA tension and requires a high protein density on the DNA as small intra-array gaps sharply decrease the blocking. These results (i) impact our mechanistic biophysical understanding of DNA loop extrusion beyond single objects on the DNA, providing a unique example of linear protein arrays that block loop extrusion with an unprecedently high efficiency, (ii) provide evidence for the hypothesis that telomere-telomere fusions preferentially break at fusion points due to a force focusing organized by Rap1-mediated condensin stalling, (iii) uncovered a new feature of telomeres and (iv) more generally highlight the intricate interplay between SMC-driven chromosomal structure, local DNA stiffness, and protein occupancy.

### Condensin is enriched at the border of Rap1 arrays

Stalling of condensin-driven DNA loop extrusion at dense telomere Rap1 arrays would result in a local accumulation of condensin at the edges of these arrays (**Fig. 1A**). To test this hypothesis, we engineered Rap1 binding-site arrays with a site density akin to native telomere sequences^12,45^ into the genome. These arrays of 16 Rap1 sites consisted of pairs of two neighboring Rap1 sites (mutually separated by 1 bp) that were separated by a constant gap that was set at either 6 or 35-bp (**Fig. 1B** and Methods). Subsequently, we used chromatin immunoprecipitation (ChIP) to map condensin-DNA interactions in the vicinity of these arrays (**Fig. 1C, Fig. S1A**). To maximize the odds of condensin encountering the Rap1-bound array, we crosslinked cells that were synchronized in late anaphase (30 minutes after release from a *cdc15-2^ts^* arrest)^12,22,44^ because condensin-dependent chromosome-arm compaction in yeast peaks in anaphase^22^.

**Figure 1.**
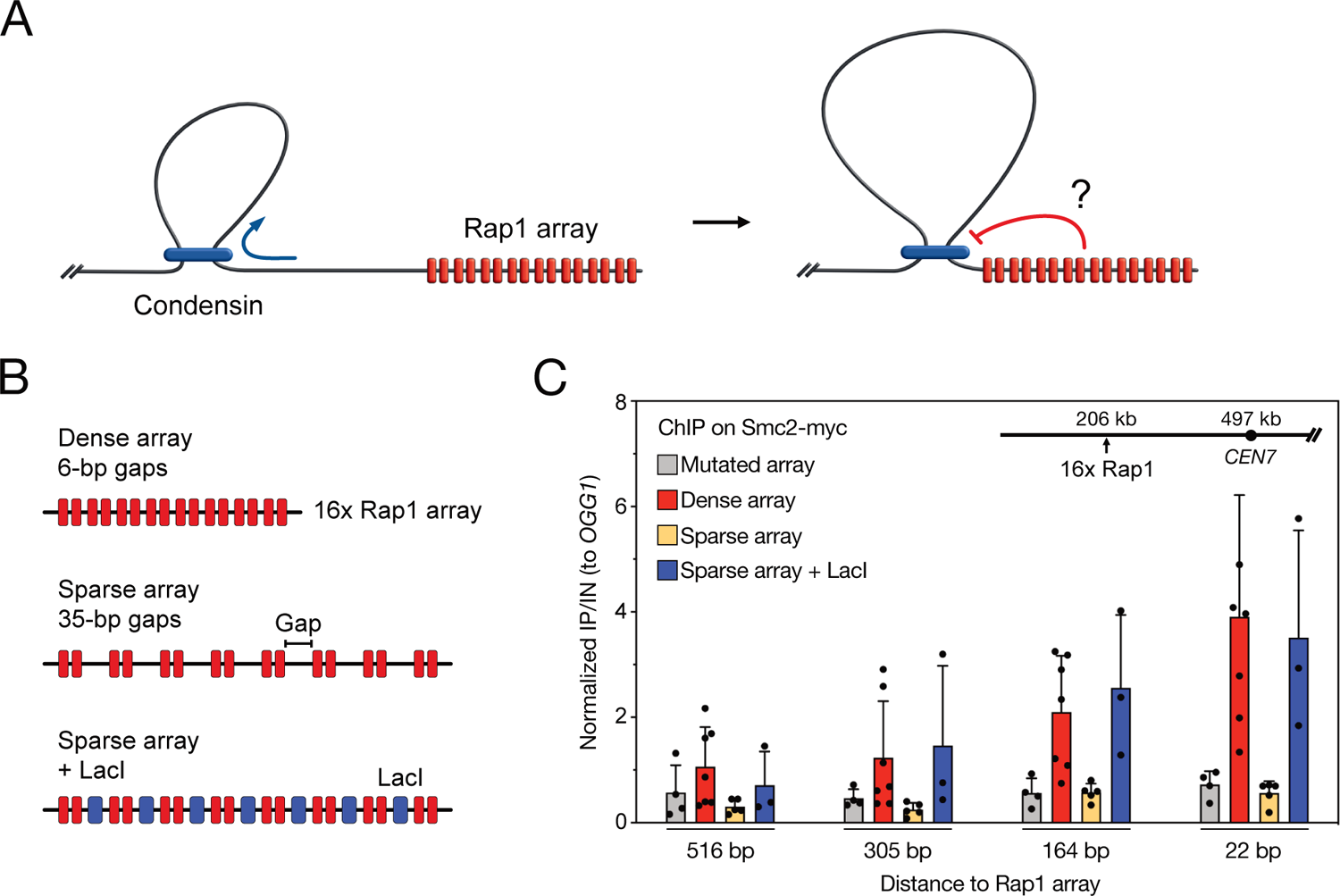
Condensin enrichment at the border of Rap1-bound arrays. **(A)** Working hypothesis for condensin stalling at a Rap1-bound array. **(B)** Scheme illustrating the Rap1 arrays used in this study. Tandem repeats are used, meaning two consecutive Rap1 binding sites separated by a gap of which we control the size. The arrays with 35-bp gaps contain a LacO sequence to which LacI protein can bind. **(C)** ChIP analysis of Smc2-Myc in cells synchronized in late anaphase. The bars represent mean IP/Input values normalized to the signal at an ectopic position; error bars indicate standard deviation over 3 or more biological replicates. Unnormalized IP/Input values are shown in Fig. S1.

We observed that an array of 16 closely spaced Rap1 sites (i.e., a dense array with 6-bp gaps) led to a 5-fold increase in the occurrence of condensin at the border of the array, relative to the level observed with an array made of mutated DNA sites that are incapable of binding Rap1^12^ (**Fig. 1C**). This condensin accumulation decreased with the distance from the array, indicating that the accumulation was most strongly localized at the edge of the Rap1 array. We saw a similar local condensin enrichment at the border of a native telomere (**Fig. S1B**). These data are in agreement with previous reports of condensin enrichment at the border of telomeres in budding yeast, fission yeast, and vertebrates during mitosis^46–48^.

If this higher condensin abundancy resulted from the stalling of condensin-driven loop extrusion at the Rap1 array, a lower density of Rap1 sites could potentially alleviate this higher abundancy, for example by exposing bare DNA segments within the array that condensin could contact during the process of loop extrusion. To investigate this, we tested an array of 16 Rap1 sites that were spaced with a 35-bp bare DNA linker between every two successive sites (**Fig. 1B**). Since Rap1 binds uncooperatively to each site^49^, such a spacer length should only impact the Rap1 density but not its high affinity^49^. As anticipated, lowering the Rap1 density strongly reduced the condensin accumulation at the border of the array (**Fig. 1C**). We conclude that condensin loop extrusion stalls at high-density Rap1 telomere arrays, but not at sparse arrays with a lower Rap1 density. Because the sparse arrays are longer than the dense arrays, this stalling is primarily due to the high local density of proteins rather than the length of the array.

To assess whether loop extrusion stalling is due to a purely physical blockade of the protein array, as opposed to possible chemical interactions, we engineered the 35-bp linker sequence to contain a *LacO* site that can be bound by lacI, thus filling the gaps between the Rap1 proteins. Notably, LacI and Rap1 bind their respective site with similar affinities^49–51^. The expression of LacI in cells harboring these 35-bp linker sequences resulted in a strongly increased abundancy of condensin at the border of the array, to the same level as the 6-bp spaced dense array (**Fig. 1C**). These findings indicate that DNA loop extrusion by condensin is stalled by the protein array due to mere physical interactions, rather than due to chemical interactions with Rap1 specifically – implying that any long dense protein array on DNA will stall condensin-driven loop extrusion.

### High-density Rap1 arrays stall loop extrusion *in vitro*

Previous *in vitro* experiments showed that, surprisingly, most single DNA-binding proteins hardly pose any barrier to loop-extruding condensin^36^. Condensin can even pass 200 nm DNA-bound beads that are larger than its ring size and accommodate those into the extruded loop^36^. Here, we used the same single-molecule-visualization assay to test whether high-density Rap1 arrays alone block loop extrusion. To this end, we inserted a Rap1 array into a long (42-kb) DNA molecule. The DNA constructs were incubated with purified and fluorescently labeled Rap1 at a 5x to 7x excess of protein to the number of Rap1 binding sites. Then, Rap1-bound DNA was flushed into a flow channel with a pegylated and biotinylated surface to which the biotinylated ends of the DNA molecules attached via biotin-streptavidin binding (**Fig. 2A**). Rap1 bound efficiently and specifically to its binding site under these conditions showing a near-100% binding efficiency and negligible off-target binding (see methods and **Fig. S2A-C**), in line with its high affinity *in vitro (K_D_* ≈ 3 nM)^49,51,52^. The residence time of Rap1 was measured under our imaging conditions (see methods and **Fig. S2D-G**), showing that Rap1 stayed bound to its binding site for much longer than our acquisition time for loop extrusion experiments (median residence time: 166 min, compared to <30 min acquisition). From these data we concluded that our linear Rap1 arrays were saturated with bound Rap1 proteins during the single-molecule experiments.

**Figure 2.**
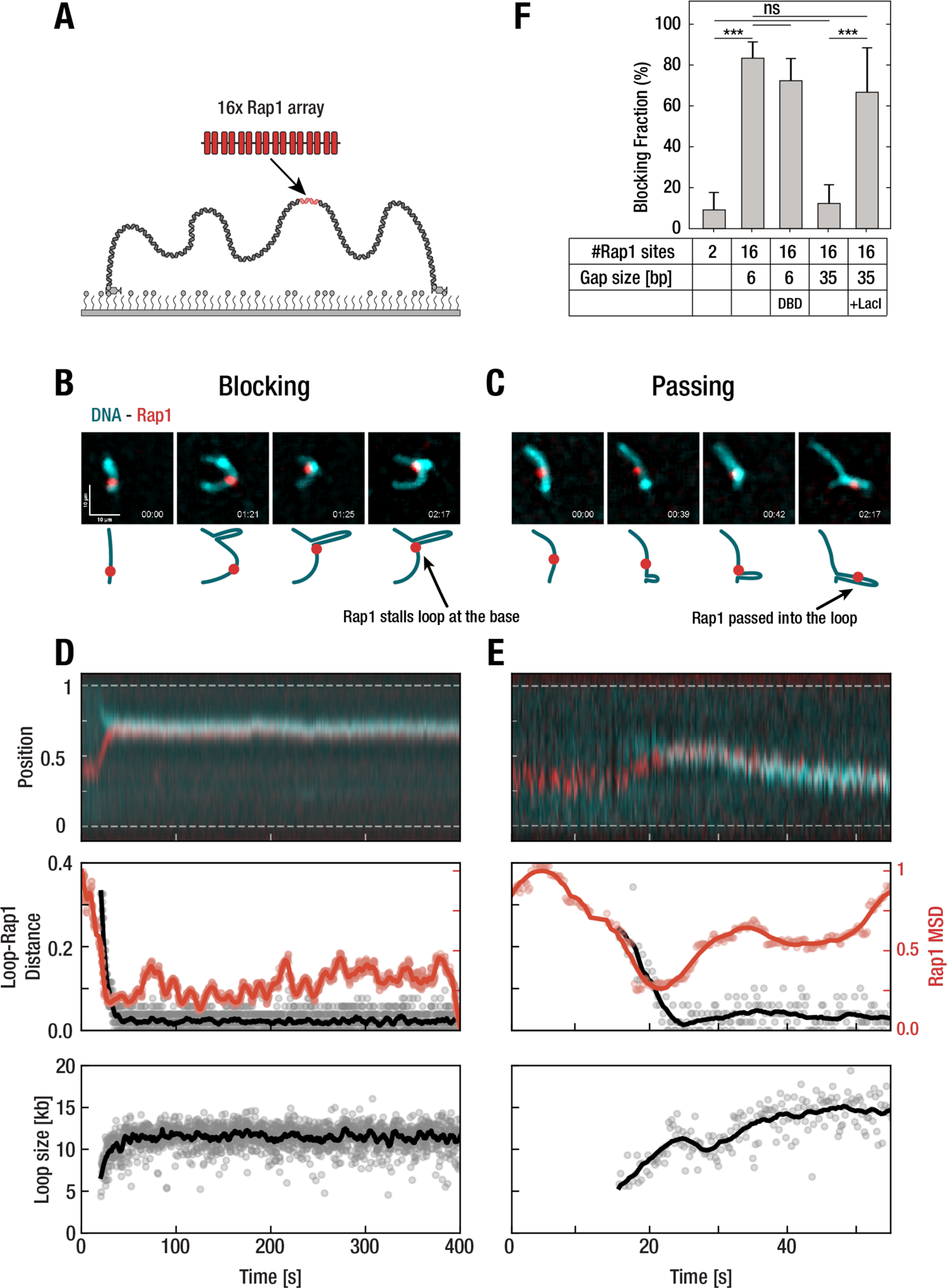
Dense Rap1 arrays are roadblocks to loop-extruding condensin in *in vitro* single-molecule experiments. **(A)** Schematic overview of a 42 kb linearized DNA molecule with a Rap1 array sequence that is tethered to a glass surface through biotin-streptavidin. **(B)** Example of a stalling event. Side-flow images of DNA in cyan and a Rap1 array red. The time points represent the initial position of Rap1 on the DNA, followed by the initiation of a loop, the encounter between the loop and Rap1, and finally the position of the Rap1 array and the loop when flow is maximal, showing that the array is at the loop base for blocking events and passes into the loop for passing events. **(C)** Same as in panel B, but for a passing event. **(D)** Kymograph analysis of a blocking event in an experiment without applied flow. Normalized positions of DNA and Rap1 are shown, as well as the distance between the loop and the Rap1 (black), and the 51-frame moving MSD of Rap1 (red). Bottom plot shows the size of the loop in kb, calculated from the fraction of fluorescence intensity of the loop relative to the whole DNA. **(E)** Same as in panel D, but for a passing event. **(F)** Blocking fraction for various Rap1 arrays. ‘DBD’ indicates the use of the DNA-binding domain truncation of Rap1 as opposed to the full-length protein. ‘+LacI’ indicates the addition of LacI protein. Error bars represent the 95% binomial confidence interval. *** indicates p<10^-3^, and *ns* indicates no significant difference as determined by the Fisher exact test.

After binding our Rap1 protein arrays in the flow cells, we next added condensin (see Methods) to observe encounters between the arrays and loop-extruding condensin. High-density linear Rap1 arrays (16 Rap1 binding sites with 6-bp gap – same as used *in vivo*) were found to clearly stall loop extrusion. This was first visualized qualitatively in a buffer flow that was applied perpendicular to the direction in which DNA was inserted. **Figure 2B** shows the typical blocking behavior where a DNA loop (cyan) developed and got stalled as soon as it encountered the Rap1 array (red); **Figure 2C** shows a passing event, where the condensin bypassed the Rap1 array and accommodated that into the extruded DNA loop. To quantify blocking in the absence of any flow (avoiding effects of the flow-associated force), imaging was performed after buffer flow was stopped. Analysis on resulting kymographs (which show the fluorescent intensity along the DNA versus time) was performed as previously described^20^. To discern stalling from passing events, we defined ‘stalling’ as an event that displayed a vanishingly small distance between the Rap1 array and the extruded loop, as well as a plateau in both the loop size and the moving mean squared displacement (MSD) (see Methods and ref. 36). By contrast, in passing events, the loop continued to grow and the moving MSD increased upon an encounter (cf. **Fig. 2B** and **Fig. 2C** for kymograph analysis). Subsequently, we estimated the blocking efficiency as the number of stalling events relative to the total number of encounters.

The high-density linear Rap1 arrays with 16 consecutively bound Rap1 proteins were found to very efficiently stall loop extrusion, with a blocking efficiency of 83 ± 8% (N=84) (**Fig. 2F**). This is an extremely high blocking efficiency, higher than measured for any other DNA-binding protein^36^, and higher than measured for encounters between cohesin and CTCF which involve chemical interactions^33^. For a block of only two Rap1 binding sites, the blocking efficiency was by contrast very low (9 ± 8%, N=44), allowing condensin-mediated loop extrusion to simply pass Rap1 into its loop in the vast majority of encounters (**Fig. 2F**). Furthermore, we found that binding of merely the Rap1 DNA-binding domain (DBD, fragment 310-608 omitting the N- and C-termini of Rap1) to the high-density 16 Rap1-site array also blocked loop-extruding condensin with a high efficiency (72 ± 11%, N=65), similar to the full-length protein (i.e., no significant difference). This shows that it is the coverage of the DNA by protein, be it Rap1 or solely its DBD, which underlies efficient blocking of loop extrusion.

By contrast, sparse Rap1 arrays with 35-bp gaps between Rap1 tandems (as in **Fig. 1**), showed a very low blocking efficiency (12 ± 9%, N=49) similar to that for only two adjacent binding sites. Inserting LacI protein into the gaps of the low-density array did, however, restore a high blocking efficiency (67 ± 22%, N=18), showing that the linear protein filament provides efficient blockage of DNA loop extrusion. In agreement with the *in vivo* findings (**Fig. 1C**), this demonstrates that stalling is primarily due to the high local density of proteins in the ∼100 nm long array. The experiments with truncated Rap1 (DBD) and with LacI insertions indicate that there is no specific protein-protein interaction between Rap1 and condensin, but instead that stalling is due to a physical rather than a biochemical interaction.

### Stalling of loop extrusion depends on array density, array length, and DNA tension

To better understand the underlying biophysical mechanism of loop extrusion stalling by the Rap1 arrays, we tested the dependence of stalling on a variety of parameters. First, we systematically examined the effects of array density on stalling. We performed our single-molecule loop-extrusion assay with Rap1 arrays that had increasingly larger gaps in between pairs of Rap1 proteins on the DNA (**Fig. 1B**). Building on our prior observation that another SMC complex, cohesin, is blocked by CTCF in a tension-dependent manner^33^, we furthermore characterized the blocking efficiency as a function of DNA tension (ranging from 0-0.2 pN) exerted on the DNA at the time of encounter. These forces are well below the stalling force of condensin, which we previously reported at ∼1 pN ^53^.

We observed an approximately linear dependence of the blocking efficiency on the gap size in all force regimes (**Fig. 3A**). While at relatively high tensions (>0.13 pN), a near-100% blocking was observed for the densest (6-bp gap) array, the blocking efficiency monotonously reduced with increasing gap sizes, to ∼10% for the 35-bp gap array. At lower DNA tension, the blocking efficiency was reduced for all arrays. The monotonous decrease of the blocking efficiency with gap size did not, within the finite signal-to noise ratio, show a clear threshold-like behavior that one would expect if there were an enabling gap size that allows condensin to make contacts within the array.

**Figure 3.**
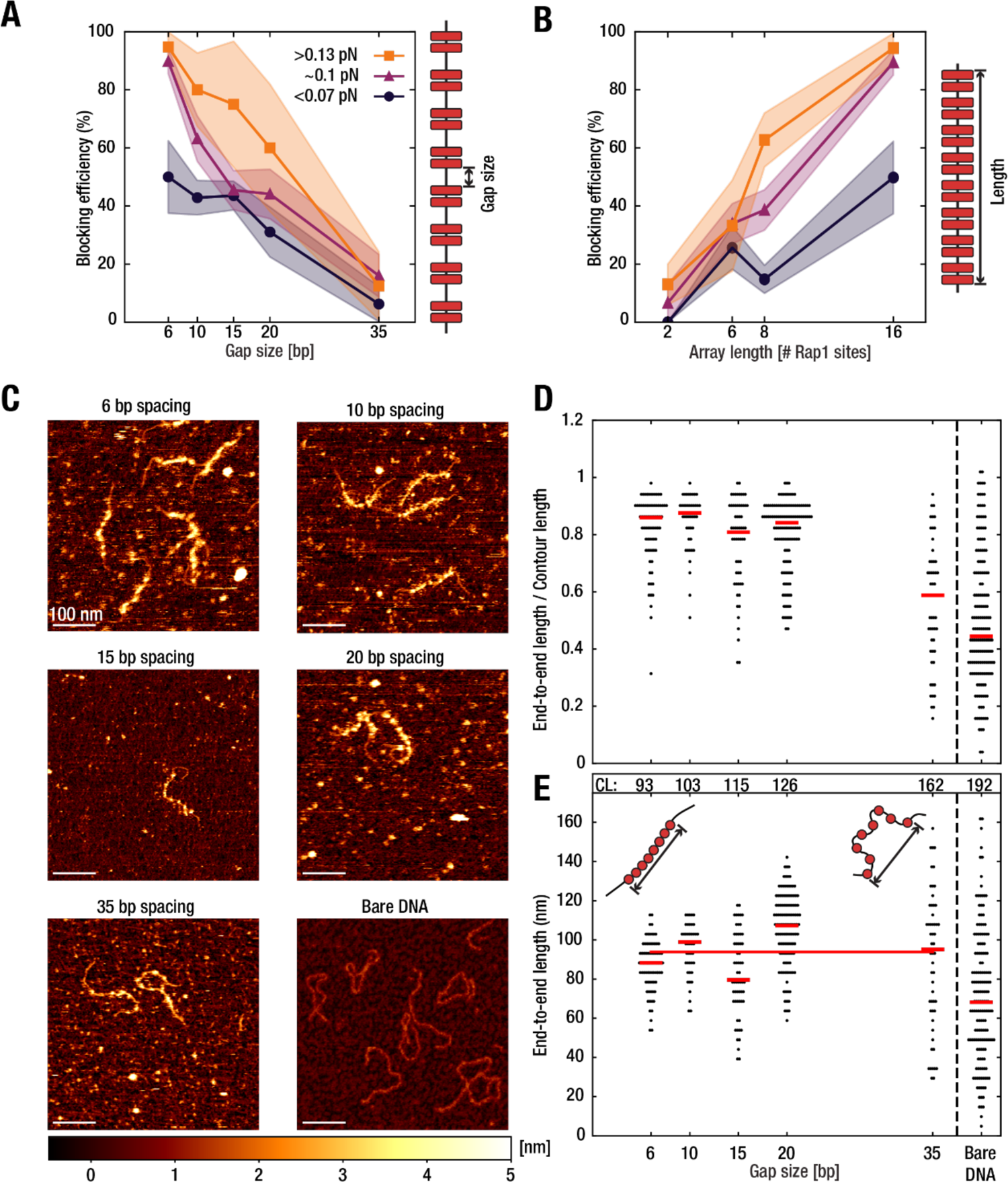
Rap1 array density and length modulate the stalling efficiency for condensin. **(A)** Blocking efficiency of Rap1 arrays with 16 binding sites versus gap size. Blocking efficiency is denoted for three force ranges. Shaded areas represent the standard error of proportion. **(B)** Blocking efficiency of Rap1 arrays versus array length, for constant 6-bp gap size. **(C)** Representative images from AFM experiments for the Rap1 arrays of panel A, and for bare DNA. **(D)** Measured end-to-end lengths normalized by the measured contour length of Rap1 arrays from AFM. **(E)** End-to-end lengths, with the contour length (CL) of each construct shown above in nm. Red bars show the median of the population. The red line shows the average end-to-end length for the different constructs (93.8 nm).

To dissect the relation between loop extrusion stalling and the length of Rap1 arrays, we next measured the blocking efficiency of dense arrays (i.e., only 6-bp gaps) with 2, 6, 8, or 16 binding sites, i.e., arrays whose length ranges from 10 to 93 nm (see **Table S2**). We observed a strong increase of the blocking efficiency with array length for all force regimes, see **Fig. 3B**, where blocking was negligible for a single Rap1 pair, but very pronounced for the 16x Rap1 array. Interestingly, the blocking efficiency exhibits a more pronounced effect of the DNA tension at higher array lengths. This suggests that local bending of DNA, which is hampered at higher DNA tension, may be important to ongoing DNA loop extrusion (refs. 18,31, and Dekker et al, Science, to appear on Nov.10).

These data show that stalling depends on array density as well as array length. Notably, in the lowest force regime, even the longest of the dense Rap1 arrays (16x Rap1 with 6-bp gaps) can still pass into the loop for a sizeable fraction (∼50%) of the encounters, which prompts us to hypothesize that condensin can occasionally grab even beyond the longest 93 nm array, in accordance with our previous measurements of the step sizes that showed that condensin occasionally makes steps larger than its ∼40 nm diameter^53,54^. The biophysical process of loop extrusion likely involves a large conformational change of the SMC complex (refs. 18,30, and Dekker et al, Science, to appear on Nov.10) as well as the polymer dynamics of the DNA (with its local Rap1 array) which is set by thermal fluctuations and polymer stiffness. Hence, we next turned to investigate how Rap1 influences the polymer properties of DNA.

To investigate how the stiffness of the Rap1 arrays depends on their density, we analyzed the structure of the arrays using atomic force microscopy (AFM). **Figure 3C** shows typical images of DNA molecules with bound Rap1 proteins, for a variety of gap sizes. The 16 Rap1 protein arrays with small (6 bp) to medium (20 bp) gaps were found to act as fairly stiff rods (in accordance with a previous report^55^) whereas the arrays became more flexible as the gap increased further to 35-bp (**Fig. 3C-E**), approaching the flexibility of bare DNA. To quantify the data, we measured the end-to-end lengths of the Rap1 arrays and normalized that to the measured contour length (**Fig. 3D**). For the densest arrays, the normalized end-to-end length approached unity, i.e., the end-to-end length thus roughly equaled the contour length, indicating that these arrays behave like stiff rods that do not bend over their length. Since the end-to-end length is only very weakly dependent on the stiffness in this length regime, quantitative conclusions about the intrinsic stiffness of the arrays cannot be drawn from these data. By contrast, the normalized end-to-end length decreased to a value of ∼0.6 for the 35-bp gap arrays with a wide distribution, indicating a greater freedom to take on different possible conformations which points to a greater flexibility. Interestingly, the mean absolute end-to-end length displayed in **Figure 3E** was found to be approximately constant with gap size. As illustrated in the insets to **Figure 3E**, this implies that the end-to-end length of the stiff 6-bp array (which equals the 93 nm contour length) happens to be about the same as the end-to-end length of the highly flexible 35-bp array which has a contour length of 162 nm. Taken together, we conclude that the denser arrays are also stiffer, which may contribute to their loop extrusion blocking efficiency.

### Condensin stalling at dense Rap1 arrays induces local chromatin decompaction in anaphase

Condensin stalling at Rap1 arrays should change local chromatin compaction in cells where condensin is active. We tested this prediction using a microscopy-based approach. We tagged two positions that were 48-kb apart on a chromosome arm with distinct LacO and TetO arrays that were bound by mCherry and GFP respectively (**Fig. 4A**). By measuring the projected 2D distance between these two spots, we inferred the local degree of chromosome folding. The median distance between the two spots decreased in cells in anaphase compared to G1 cells. This chromosome compaction did, however, not occur in condensin-depleted cells (**Fig. 4A**, *smc2-AID* +IAA), indicating that it resulted from condensin activity during anaphase^22,56–59^.

**Figure 4.**
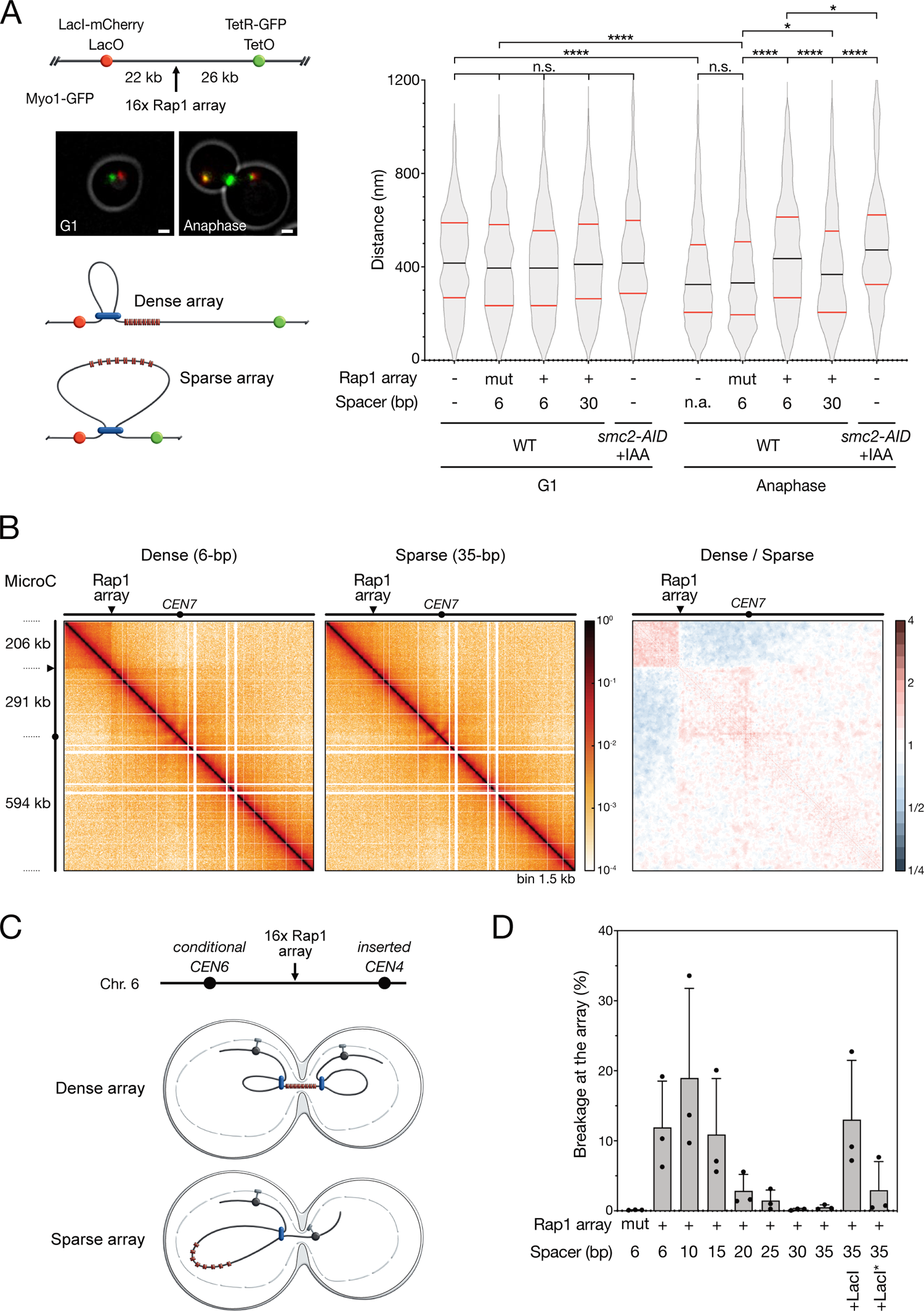
Impact of Rap1-bound array on chromatin compaction in anaphase. **(A)** A telomere-like Rap1 array causes local chromatin decompaction in anaphase. Representative images of cells at distinct cell cycle stages. In anaphase, the two sister chromatids are separated and the actomyosin ring (Myo1-GFP labelled) is open. In G1 cells, the ring is disassembled. Scale bars: 1 µm (pixel size: 65 nm). Distance between the two fluorescent spots shown in G1 cells and in anaphase cells. Black lines indicate median value, red lines quartiles value. Statistical significance is given by the Mann-Whitney test. **(B)** MicroC contact maps of chromosome 7 in cells synchronized in late anaphase. The ratios were determined using the Serpentine tool^82^. Whole-genome contact maps are shown in Fig. S3. **(C)** Condensin stalling at dense Rap1 arrays in dicentrics focuses entrapment and breakage by abscission. **(D)** Dicentric breakage at Rap1-bound arrays with varying gap sizes. The error bars represent the standard deviation over 3 biological replicates. Gels for each single experiment are shown in Fig. S4.

The insertion of a dense array (16 Rap1 sites with 6-bp gap) half-way between the two fluorescently tagged positions was found to have no impact in G1 cells (**Fig. 4A**), which was expected given the low condensin activity. It also indicates that the local DNA stiffening of the array had no impact on the chromosome compaction at this scale. In cells in anaphase, however, inclusion of the dense array increased the median distance between the two spots to ∼400 nm, equivalent to the low compaction seen in G1. This indicates that a Rap1-bound array caused a local chromatin decompaction during anaphase. Lowering the Rap1 density of the array restored the anaphase compaction (cf. 30-bp gap in **Fig. 4A**). The sensitivity to Rap1 density shows that it is the condensin stalling at the array that causes the observed chromatin decompaction by preventing the formation of larger loops that brings the two spots in closer proximity.

To further validate this result, we used a MicroC approach^60,61^ to quantify the frequency of contacts between adjacent chromatin regions in cells synchronized in late anaphase. Strikingly, a dense Rap1 array (16 Rap1 sites with 6-bp gaps) reduced the contacts of the telomere-proximal chromosome region with the rest of the chromosome arm (**Fig. 4B**, **Fig. S3**), the expected outcome of condensin stalling at the dense array. A sparse array (35-bp gaps, **Fig. 4B**, **Fig. S3**) and an array of mutated DNA sites incapable of binding Rap1 (**Fig. S3**) failed to insulate the telomere-proximal region, in accordance with condensin being able to extrude these arrays in an unhindered way. This shows that the reduced contact frequency caused by the dense array stems from a reduced frequency of loops that would bring DNA together from the two sides separated by the array.

Preferential breakage of dicentric chromosomes near Rap1 arrays is another anticipated outcome of condensin stalling at the arrays (**Fig. 4C**)^12^. We used this readout to test arrays with various Rap1 densities. Rap1 arrays of 16 binding sites with gaps ranging from 6 to 35-bp were inserted in a conditional dicentric chromosome, whose one centromere can be reversibly inactivated (**Fig. 4C**). To monitor dicentric breakage by abscission, we reactivated the conditional centromere in cells synchronously released from a G1 arrest. Cells were harvested either prior to dicentric breakage (nocodazole arrest) or after dicentric breakage in the next G1 (alpha factor arrest). Chromosome fragments were separated by Pulse Field Gel Electrophoresis (PFGE) and detected by Southern blot. In the absence of Rap1 arrays between centromeres, dicentric breakage preferentially occurred near the centromeres (**Fig. S4**)^12,44^.

We observed a strong dependence of dicentric breakage on the gap size. Only high-density arrays focused the breakage at the array, while low-density arrays with gaps of 30 and 35-bp failed to do so (**Fig. 4D**). Ectopic expression of bacterial LacI restored a strong breakage at the arrays with the 35-bp gap sequence containing a *LacO* site, as reported previously^12^. This effect was attenuated when utilizing a LacI* allele with a reduced *LacO* affinity (**Fig. 4D**)^62^. These *in vivo* results further indicate that continuous high-affinity protein binding along the array on DNA is a key feature needed to stall condensin.

## DISCUSSION

This work shows that DNA coverage by a telomere protein strongly modulates condensin loop extrusion activity *in vivo* and *in vitro*. DNA-loop-extruding condensin stalls at encounters with telomere-like arrays of Rap1 protein bound on DNA in a length- and density-dependent manner. While individual DNA-bound roadblocks can easily pass into the extruded loop^36^, a dense coverage of DNA by proteins halts loop extrusion. Such a stalling of loop extrusion results in a local boundary to chromosomal compaction during anaphase. Notably, these telomeric protein arrays exhibit a remarkable stability as the Rap1 residence time on DNA is of the order of hours (**Fig. S2E**), which is much longer than the inverse stepping rate of the loop extrusion.

Our observations have implications for our biophysical understanding of loop extrusion by SMC complexes. Rap1 binding into a closely spaced array that covers the DNA makes this region inaccessible to a loop extruder. We found that small gaps in the array facilitate passage, and larger gaps of ∼30-bp even allow unhindered loop extrusion through the array (**Fig. 1C**, **Fig. 2F**, **Fig. 3A**, **Fig. 4**). While the data thus clearly point to a steric hindrance effect where Rap1 precludes the availability of DNA as a substrate for loop extrusion, it is of interest to ask whether the increased local stiffness plays a role as well, since Rap1 binding stiffens the DNA. Such a stiffening can potentially hinder loop extrusion in two ways. First, it may be energetically costly to reel the new DNA within the SMC lumen due to its reduced flexibility, as current models for loop extrusion predict a significant bending of DNA during a loop extrusion step (refs. 18,53,54,63–66 and Dekker et al, Science, to appear on Nov.10). Second, a stiffer Rap1 array positions the next freely accessible DNA further away from condensin, making it less likely that the SMC can reach beyond the array. While we observed differences between Rap1 arrays of varying length and density, it is difficult to disentangle the effects of density and stiffness, and therefore we cannot unambiguously determine the relative importance of the stiffness. The data call for a detailed mechanistic model and simulations of loop extrusion of DNA with a local array of varying stiffness. Summing up, we conclude that the dense linear protein arrays stall condensin by reducing the amount of freely accessible DNA that can be grabbed and processed by condensin, as well as potentially by inhibiting the incorporation of the array into the loop and by distancing the freely accessible DNA to positions beyond the array.

Condensin stalling by Rap1 at telomere-telomere fusions favors dicentric breakage near the fusion points. This mechanism provides a back-up for telomere protection and contributes to genome stability^42^. As corroborated by microscopy and MicroC analyses (**Fig. 4A&B**), we find that dense Rap1 arrays establish boundaries to loop extrusion during anaphase, resulting in local chromatin insulation. This reveals a mechanism underlying the positioning of dicentric breakage at telomere-telomere fusions. In anaphase, the connection of centromeres to the spindle poles stretches dicentric anaphase bridges. As telophase progresses, the disassembly of the mitotic spindle and the detachment of the spindle poles from the cell cortex allow condensin to recoil the dicentric bridges prior to septum closure^12,44^. Condensin stalling at telomere-telomere fusions will favor the creation of two distinct domains, one in each nuclear lobe, out of the two chromosome regions that are separated by the fusion point. This spatial insulation will direct the telomere-telomere fusion toward the midzone, where the septum grows, thus resulting in its entrapment and breakage by abscission.

Our findings show that the repeated nature of telomeres and the consequential dense DNA coverage yield a unique 1D property: the ability to inhibit protein machines acting along the DNA. The blocking of SMC-driven loop extrusion could apply more broadly to other activities whose control is important to telomere functions. Apart from its role in resolving dicentric chromosomes, it is conceivable that condensin stalling at native unfused telomeres contributes to their accurate segregation (**Fig. S5**). Without such stalling, loop extrusion would proceed unhindered until the end of chromosomes, where condensin would run off the DNA, leaving the chromosome ends uncompacted. Instead, a stalling of loop extrusion at the chromosome ends ensures their individualization and proper compaction, facilitating their correct segregation prior to cell division. In this way, condensin stalling at telomeres might further contribute to genome stability.

As we found that extended linear protein filaments can stall condensin-driven loop extrusion remarkably efficiently, it is of interest to ask whether this result can be generalized, i.e., whether linear protein filaments more generally block SMCs to extrude loops of DNA. Several observations indicate that this indeed may be the case. DNA repair of double-stranded breaks (DSBs) features a stage where DNA is coated with dense protein arrays, and it was reported that cohesin accumulates at these filaments^67,68^. While it is commonly assumed that cohesin is specifically loaded at DSB sites^68^, loop extrusion could play a role in targeting cohesin to these sites^69,70^. Furthermore, it was shown that highly transcribed genes significantly slow down loop-extruding SMC complexes^32,39,40^. Possibly this can be attributed to a local dense coverage of DNA by RNA-polymerases that line up in long ‘trains’. Finally, as the linker length needed for loop extrusion through Rap1 arrays approximates the average spacing between nucleosomes^71^, it will be of interest to see if dense nucleosome fibers block SMCs. The tension that condensin can exert on chromatin (<1 pN^15^) is insufficient to unwrap nucleosomes^72^ but may be sufficient to stretch them^73,74^, which could help to expose the internucleosomal DNA for capture by the SMC complex during loop extrusion, a hypothesis that remains to be tested.

Loop extrusion stands as a universally conserved mechanism across the SMC family (refs. 15–17,19,20 and Dekker et al, Science, to appear on Nov.10). While we presented a detailed study of condensin and Rap1 in *S. cerevisiae,* we estimate that our findings have a general significance and likely also hold for other SMCs and other protein filaments – providing an important control element for chromosome organization.

## MATERIALS & METHODS

### Strains

All yeast strains used in this study are listed in **Table S1**.

### Cell cycle synchronization

To synchronized cells in late anaphase (ChIP and MicroC experiments), exponentially growing cells carrying the *cdc15-2* thermosensitive allele were arrested at restrictive temperature (36°C) for about 90 minutes prior to be shifted back at permissive temperature (25°C) for 30 minutes. To assess dicentric breakage, cells growing exponentially in galactose-containing synthetic medium (*CEN6* OFF) were arrested in G1 with α-factor (10^−7^ M). Cells were released from the G1 arrest with two washes in glucose-containing rich medium (YPD). Half the culture was complemented with nocodazole (5 μg/mL) to arrest the cells in G2/M. The other half was complemented with α-factor (10^−7^ M) about one hour after the washes to arrest the cells in the next G1.

### Pulse-Field Gel Electrophoretic (PFGE) and Southern blot

Yeast DNA embedded in agarose plugs was prepared as described ^12^ with minor modification (see supplementary information). Pulse-field gel electrophoresis was carried out in a 0.9% agarose gel in 0.5× TBE at 14°C with a CHEF DR III from Bio-Rad with a constant switch time of 20 s during 24 h. Gel-Red labeled DNA was detected by a Typhoon scanner. DNA transferred to a nitrocellulose member was hybridized with ^32^P-labeled *TUB2* (chr. 6 probe) and *POL4* (chr. 3 probe) fragment as previously described ^12^.

### Distance measurements by microscopy of cells

Exponential growing cells (0.8 OD) in rich medium (YPD) were washed in synthetic medium prior to live-cell imaging with a wide-field inverted microscope (Leica DMI-6000B) equipped with Adaptive Focus Control to eliminate Z drift, a 100×/1.4 NA immersion objective with a Prior NanoScanZ Nanopositioning Piezo Z Stage System, a CMOS camera (ORCA-Flash4.0; Hamamatsu), and a solid-state light source (SpectraX, Lumencore). The system is piloted by MetaMorph software (Molecular Device). 2mM Indole-3-acetic acid (IAA) was added to exponential growing *smc2-AID* cells in YPD for 1 hour prior to imaging.

GFP and mCherry two-color images were acquired over 19 focal steps of 0.2µm using solid state 475 and 575nm diodes and appropriate filters (GFP-mRFP filter; excitation: double BP, 450–490/550– 590nm and dichroic double BP 500–550/600–665nm; Chroma Technology Corp.). Acquisition of both wavelengths was completed on each focal plane with an exposure time of 50ms, before 0.2µm steps, to minimise the possibility of array movement between acquisitions of each wavelength. A single bright-field image on one focal plane was acquired at each time point with an exposure of 50ms. All images shown are maximum intensity z projections of z-stack images.

Image analysis was achieved following processing with ImageJ Fiji software, using scripts written in ImageJ macro language. Briefly, local maxima that define GFP and mCherry fluorescent array positions were determined from 2D maximal projections of three-dimensional data sets. Fluorescent signals within cells were confirmed manually from 3 color merged images. The distance between the two closest GFP and mCherry maxima was calculated using their extracted XY coordinates in R software (v4.1.1).

### ChIP

ChIP experiments were carried out as previously described with minor modifications ^75,76^.

### MicroC-XL

Micro-C was done following a mixed protocol described previously^61,77^ with minor modification. Briefly, 55 OD anaphase blocked yeast cultures were crosslinked with 3% formaldehyde for 15 min at 30°C. The reactions were quenched with 250 mM glycine at 30°C temperature for 5 min with agitation. Cells were pelleted by centrifugation at 4000 rpm at 4°C for 5 min and washed twice with water. Cells were then resuspended in Buffer Z (1M sorbitol, 50 mM Tris-HCl pH 7.4, 10 mM β-mercaptoethanol) and spheroplasted by addition of 250 ug/mL Zymolyase (MP08320932) at 30°C in an incubator at 200 rpm for 40 to 60 minutes. Spheroplasts were washed once by 4°C PBS and then pelleted at 4000 rpm at 4°C for 10 min. Pellets were re-crosslinked by addition of PBS supplemented with 3 mM disuccinimidyl glutarate (ThermoFisher #20593) and incubated at 30°C for 40 min with gentle shaking before quenching by addition of 400 mM final glycine for 5 minutes at 30°C. Cells were pelleted by centrifugation at 4000 rpm at 4°C for 10 min, washed once with ice-cold PBS and stored at −80°C. Pellets were treated as previously described^61^ up to the decrosslink part. Decrosslink solution was added with an equal volume of Phenol:Chloroform:Isoamyl Alcohol (25:24:1), vortexed intensively centrifuged for 15 minutes at room temperature. The aqueous phase loaded and purified on ZymoClean column according to the manufacturer protocol. Dinucleosomes were purified and excised from a 3% NuSieve GTG agarose gel (Lonza #50081) using Zymoclean Gel DNA Recovery Kit (Zymo #D4008). Micro-C libraries were prepared using the NEBNext Ultra II DNA Library Prep Kit for Illumina (NEB #E7645) following^77^ manufacturer instructions and sequenced on the Illumina NovaSeq 6000 platform.

Micro-C datasets were analysed using the Distiller pipeline (https://github.com/open2c/distiller, commit 8aa86e) to implement read filtering, alignment, PCR duplicate removal, and binning and balancing of replicate and sample matrices. Reads were aligned to W303 using bwa 0.17.7 and the resulting maps filtered to remove low-quality alignments (MAPQ<30) and cis alignment pairs within 150 bp. Replicates were analysed independently, and their quality assessed before aggregation into sample-level datasets. Maps were visualized and explored using Higlass^78^.

### DNA preparation for single-molecule-visualization assay

42-kb Linear cosmid-i95 plasmids with inserted sequences were prepared as previously reported^36,79^. First, the i95-cosmid was linearized with PsiI-v2 (New England Biolabs). Second, the remaining 5’-phosphate groups were dephosphorylated using calf-intestinal alkaline phosphatase for 10 minutes at 37°C and finally heat inactivated for 20 min at 80°C (Quick CIP, New England Biolabs). The Rap1 arrays initially cloned in a pUC19-derived vector (**Table S2**) were digested with PvuII (New England Biolabs) and subsequently gel isolated. The fragments were ligated together by using a T4 DNA ligase in T4 ligase buffer (New England Biolabs), with 1 mM ATP overnight at 16°C. The final constructs were transformed into *E. coli* NEB 10-beta cells (New England Biolabs) and all constructs were sequence verified using plasmidsaurus Oxford Nanopore long read sequencing. Inserted sequences are listed in **Table S2**. To linearize these cosmids and prepare them for flow cell insertion, the cosmids were isolated using a Midiprep and a QIAfilter plasmid midi kit (QIAGEN). The cosmids were then digested for 2 hours at 37°C and heat-inactivated for 20 minutes at 80°C using SpeI-HF (New England Biolabs). Next, 5’-biotin handles were constructed by a PCR reaction from a pBluescript SK+ (StrataGene) using 5’-biotin primers JT337 (Bio-AGAATAGACCGAGATAGGGTTGAGTG) and JT338 (Bio-GGCAGGGTCGGAACAGGAGAG). The PCR fragment was then digested by the same procedure as for the large cosmid, resulting in ∼600-bp 5’-biotin handles, which were mixed with the digested cosmids in a 10x excess before ligation by T4 DNA ligase in T4 ligase buffer (New England Biolabs) at 16°C overnight. The reaction was subsequently heat-inactivated at 65°C for 25 min. The final linear construct was cleanup using an ÄKTA Start (Cytiva), with a homemade gel filtration column containing 46 mL of Sephacryl S-1000 SF gel filtration media, run with TE + 150 mM NaCl buffer at 0.5 mL/min.

### Protein purification and fluorescent labelling

His6-TEV-4G-ScRap1-1-827 (Rap1 full length) was induced with 0.5 mM isopropyl-β-D-thiogalactoside (IPTG) four hours at 30 °C into *E. coli* strain BL21 (DE3) STAR (Invitrogen). All of the subsequent protein purification steps were carried out at 4 °C. Cells were harvested, suspended in lysis buffer (50 mM Tris HCl [pH8@4 °C], 1M NaCl, 1 mM DTT, 20 mM Imidazole 1 mg/mL lysozyme, 1 mM 4-(2-aminoethyl) benzenesulphonyl fluoride (AEBSF), 10 mM benzaminide, 2 µM pepstatin) and disrupted by sonication. Extract was cleared by centrifugation at 186,000*g* for 1 hour at 4 °C and then incubated at 4 °C with NiNTA resin (QIAGEN) for 4 h. Mixture was poured into an Econo-Column^®^Chromatography column (BIO-RAD). After extensive washing of the resin first with buffer A (20 mM Tris HCl [pH8@4 °C], 500 mM NaCl, 1 mM DTT, 20 mM Imidazole) and then with buffer B (20 mM Tris HCl [pH8@4 °C], 100 mM NaCl, 1 mM DTT, 40 mM Imidazole), protein was eluted with buffer B complemented with 400 mM imidazole. Fractions containing purified His6-TEV-4G-ScRap1-1-827 were pooled and applied to a ResourceQ 1ml column (Cytiva) equilibrated with buffer C (20 mM Tris HCl [pH8@4 °C], 100 mM NaCl, 1 mM DTT, 1mM EDTA). Protein was eluted with a 20 mL linear gradient of 0.1–1 M NaCl. Fractions containing the purified protein were pooled and directly applied to a 1 ml HiTrap Heparin HP column (Cytiva) equilibrated with buffer C. A 30 mL linear gradient of 0.1-0.8 M NaCl was performed. TEV protease was added to the pooled fractions containing purified His6-TEV-4G-ScRap1-1-827 and the mixture was directly dialyzed against buffer D (20 mM Tris HCl [pH8@4 °C], 150 mM NaCl, 1 mM DTT, 1mM EDTA) at 4°C overnight. The mixture was then incubated with NiNTA resin (QIAGEN) for 2 hours and the purified 4G-ScRap1-1-827 without its His6-TEV tag was recovered into the flow trough. Concentration was determined using Bradford protein assay with BSA as standard. 4G-ScRap1-310-608 (Rap1 DBD) was purified with the same protocol except the HiTrap Heparin HP column which was omitted.

Rap1 protein was subsequently labelled with Janelia Fluor 646 (JF646) using a sortase reaction followed by ÄKTA purification in a MonoQ column against a 1M NaCl gradient. The labelling efficiency of Rap1-JF646 was estimated to be about 70% from the fluorophore and protein concentrations. *S. cerevisiae* condensin was purified as described in Ganji et al.^15^.

### Single-molecule-visualization assay

For the single-molecule loop extrusion assay, flow cells were prepared as previously reported^15^. Briefly, glass slides and coverslips were cleaned using successive rounds of sonication in acetone and 1M KOH followed by piranha etching. The glass surface was functionalised using aminosalinization and the surface was passivated using mPEG-SVA (Laysan Bio) and MS(PEG)_4_-NHS-Ester (Laysan Bio) in the presence of biotin-PEG-SVA (Laysan Bio). Before experiments, the flowcell was briefly incubated with streptavidin (MP Biomedicals) in T20 buffer (40 mM tris-HCl pH 8.0, 20 mM NaCl, 0.2 mM EDTA) and with 5 mg/ml BSA (ThermoFisher Scientific) also in T20 buffer. Rap1 was bound to the long linear constructs by incubating at room temperature at a 5-fold excess of protein to binding site for at least 1h in 100 mM KGlu, 2.5 mM MgCl_2_, 20 mM Tris pH 7.4, 1 mM DTT, 0.25 mg/ml BSA. After incubation, the Rap1-DNA complex was flushed into the flowcell. DNA was visualized by adding 100 nM SytoxOrange (SxO) DNA dye. Unbound complexes were flushed out and the buffer was changed to loop extrusion/imaging buffer (50 mM KGlu, 2.5 mM MgCl_2_, 40 mM Tris pH 7.5, 2 mM Trolox, 1 mM DTT, 0.25 mg/ml BSA, 5% glucose, 10 nM catalase, 18.75 nM glucose oxidase, 2 mM ATP). Purified yeast condensin was added at 2 nM in imaging buffer at a flow rate of 0.5 μL/min until loops were observed and the flow was stopped. Imaging was done with a HILO microscope, as previously described^15^, with a red (637 nm, 15 mW) and a green laser (561 nm, 0.2 mW) in alternating light excitation mode.

### Rap1-DNA binding efficiency, specificity, and residence time

Binding efficiency was estimated to be near-100% from fluorophore bleaching in our single-molecule fluorescence visualization assay. To this end, Rap1-JF646 was incubated with linear constructs that contain 2 tandem Rap1 binding sites. After flushing the binding reaction into the flow cell, bleaching was done at 25mW power with the 637 nm laser. Individual fluorescent spots were tracked and their fluorescence plotted as in **Fig. S2A**. The number of observed bleaching steps were counted for 46 spots using a step-finding hidden Markov model (sfHMM)^80^. As shown in **Fig. S2B**, half of the traces showed 1 and the other half showed 2 bleaching steps (50 ±14%, N=46). This distribution of bleaching steps is in good accordance with the 70% labelling efficiency, from which we expect to observe 54% of traces with a Rap1-JF646 signal to show two steps if both binding sites are occupied (E^2^/(E^2^ + 2E(1-E)), where E is the labelling efficiency). There is no significant difference between the expected 54% and our observed 50% (p=0.16, one-sided binomial test), indicating a near-100% binding efficiency. Binding specificity was visualized using the binding positions along the DNA molecule from the same bleaching experiment (**Fig. S2C**). The Rap1 binding sites were positioned at roughly 40% along the DNA.

To measure the residence time (**Fig. S2D-G**) of Rap1 to its binding site, we used an assay similar to that described above for determining the binding specificity. Briefly, we used a DNA construct with 2 tandem Rap1 binding sites, incubated with Rap1 at a ratio of protein to binding site of 10, for >1h at room temperature. Imaging was performed in the imaging buffer described above without ATP with an oxygen scavenger system to minimize photobleaching (50 mM KGlu, 2.5 mM MgCl_2_, 40 mM Tris pH 7.5, 2 mM Trolox, 1 mM DTT, 0.25 mg/ml BSA, 5% glucose, 10 nM catalase, 18.75 nM glucose oxidase). The constructs were imaged for 3h with infrequent imaging (1 image per 4s) to reduce photobleaching for this long measurement. Data was analyzed similar to described above, where unbinding events were counted as a downward step in kymographs, and steps were analyzed using sfHMM^80^.

### Atomic force microscopy

For AFM imaging, short DNA fragments containing Rap1 binding site arrays were produced. The same Rap1 arrays as for the single-molecule loop extrusion were cut with PvuII (New England Biolabs) and fragments containing Rap1 repeats were separated from the backbone using a similar ÄKTA procedure as mentioned above: ÄKTA Start (Cytiva) with a homemade gel filtration column containing 46 mL of Sephacryl S-1000 SF gel filtration media, run with TE + 150 mM NaCl buffer at 0.5 mL/min. To concentrate the sample, we used the vacufuge plus speedvac (Eppendorf) to reduce the volume. Next, the Rap1 array fragments were dialyzed to water to remove excess salt. Final concentration of DNA fragments was between 1 and 14 nM.

Samples were prepared by mixing DNA at a concentration of 0.5 nM with Rap1 to a protein:binding site ratio of 4.1. We used the same fluorescently labelled full-length Rap1 proteins as in the single-molecule visualization assay. Samples were incubated in a similar buffer as for the single-molecule visualization assay: 100 mM KGlu, 20 mM Tris pH 7.4, 1 mM DTT. Some of the samples had slightly higher ratios of protein to binding site due to a calculation error, but we found no significant effect on the binding in this concentration regime.

For surface deposition and measurement, we prepared mica substrates by punching 3.2 mm mica discs from mica sheets (V4 grade, SPI supplies) and gluing them to magnetic stainless steel discs with 2-component epoxy glue. The mica discs were cleaved with adhesive tape before each preparation to provide a clean surface. To ensure stable adhesion of the DNA to the mica surface, poly-L-lysine (PLL) was deposited onto the mica at a concentration of 0.01% (w/v), incubated for 3 minutes, washed with pure water and dried in a stream of nitrogen. We found that shorter incubation of the PLL, as well as the use of poly-L-orthinine instead of PLL, would lead to incomplete coverage of the mica, which promoted alignment of parts of the DNA molecules along straight lines separated by angles of 60 degrees, presumably parallel to the crystal axes of the mica. DNA-Rap1 samples were incubated for 45-90 minutes at room temperature (21°C) and then 3 µl drops were deposited onto the PLL-coated mica substrates. After 1 minute, the sample was gently washed using 200 µl of buffer applied and extracted with two separate pipettes. The sample was then placed onto the microscope and imaged in buffer. The microscope was a Bruker Multimode, with NanoScope V controller and version 9.1 Nanoscope software. The imaging mode was PeakForce QNM, with a tapping frequency of 4 kHz and a force setpoint and amplitude manually tuned for optimal image quality, typically 100 pN and 12 nm. Images were acquired with a pixel size of 2 nm, and processed for subtraction of background artifacts using the ‘align rows’ and ‘remove polynomial background’ filters in Gwyddion^81^.

To analyze the contour lengths and array end-to-end lengths, we used a homebuilt Matlab analysis package named DNAcontour. This package in the version that was used to produce the data presented here, is available at https://gitlab.tudelft.nl/allards-matlab-repo/allards-matlab-repo/-/tree/RAP1_paper/DNAcontour. To automatically select DNA molecules from AFM images, we first applied smoothing using a multi-pass Gaussian blur, followed by thresholding and filtering based on a potential DNA molecule’s height, size, and aspect ratio. To calculate the trajectories of the DNA molecules, they were skeletonized and resulting branches were connected. The initial guess for trajectories was obtained by connecting the branches with Dijkstra’s algorithm and taking the longest shortest path between endpoints. These trajectories were manually corrected where needed and the start- and endpoints of the Rap1 arrays were manually annotated. The obtained trajectories were then iteratively refined by optimizing the trajectories for the local maxima in a smooth curve. The end-to-end lengths were then calculated as the distance between the endpoints of the Rap1 arrays and the contour length was determined from the length of the DNA trajectories as shown in Fig. 3D.

## ACKNOWLEDGMENTS

We thank Milos Tisma, Theo Koenig, Eric Le Cam, Olivier Pietrement, Gerard Mazón, Armelle Lengronne, Jonathan Heuze, Olivier Alibert, Stéphane Coulon, Sylvie Tournier, Pascal Bernard, Frédéric Beckouet, Sabrina Pobieaga and Florian Roisné-Hamelin for discussions, Anders Barth for help with determining the Rap1 labelling efficiency, and Eli van der Sluis and Ashmiani van den Berg for purifying condensin and LacI proteins. Research in the laboratory of S.M. was supported by grants from *Agence Nationale de la Recherche* (ANR-14-CE10-021 DICENS, ANR-15CE12-0007 DNA-Life), *Fondation ARC pour la Recherche sur le Cancer*, *Ligue contre le Cancer*, CEA Radiation biology program and GGP CEA EDF program. A.D. was supported by a PhD fellowship from CEA and a *Ligue contre le Cancer* young researcher grant. Research in the laboratory of C.D. was supported by ERC Advanced Grant 883684 (DNA looping) NWO grant OCENW.GROOT.2019.012, and the BaSyC program.

