## Supplemental Information for "Telomere protein arrays stall DNA loop extrusion by condensin"

**Table S1: Yeast strains used in this study.**

| Name | Genotype | Figures |
|---|---|---|
| YCB212 | *MATα cdc15-2ts smc2-AID-9myc::HPHBr NATr::16 Rap1 sites 6bp gaps in 7L at position 206707* | *1C, S1* |
| YCB214 | *MATα cdc15-2ts NATr::16 Rap1 sites 6bp gaps in 7L at position 206707* | *1C, S1, 4B* |
| YCB215 | *MATα cdc15-2ts* | *4B* |
| YCB257 | *MATα cdc15-2ts smc2-AID-9myc::HPHBr NATr::16 mutated Rap1 sites 6bp gaps in 7L at position 206707* | *1C, S1* |
| YCB261 | *MATα cdc15-2ts NATr::16 mutated Rap1 sites 6bp gaps in 7L at position 206707* | *1C, S1, 4B* |
| YCB542 | *MATα cdc15-2ts smc2-AID-9myc::HPHBr NATr::16 Rap1 sites 35b gaps in 7L at position 206707* | *1C, S1* |
| YCB544 | *MATα cdc15-2ts NATr::16 Rap1 sites 35bp gaps in 7L at position 206707* | *4B* |
| YCB550 | *MATα cdc15-2ts NATr::14 Rap1 sites 35bp gaps in 7L at position 206707* | *1C, S1* |
| YCB551 | *MATα cdc15-2ts smc2-AID-9myc::HPHBr NATr::16 Rap1 sites 35bp gaps in 7L at position 206707 ura3-1::lacI-mCherry::URA3* | *1C, S1* |
| YCB552 | *MATα cdc15-2ts smc2-AID-9myc::HPHBr NATr::16 Rap1 sites 35bp gaps in 7L at position 206707 ura3-1::URA3* | *1C, S1* |
| YCB554 | *MATα cdc15-2ts NATr::16 Rap1 sites 35bp gaps in 7L at position 206707 ura3-1::URA3* | *1C, S1* |
| YCB557 | *MATα cdc15-2ts NATr::16 Rap1 sites 35bp gaps in 7L at position 206707 ura3-1::lacI-mCherry::URA3* | *1C, S1* |
| YAD82 | *MATa bar1-Δ KANr-pGAL1-CEN6-pGAL1-skHIS3 ADE2 NATr::16 mutated Rap1 sites 25bp gaps in chromosome 6 at position 192140 CEN4::klLEU2 in chromosome 6 at position 245710* | S4 |
| YAD83 | *MATa bar1-Δ KANr-pGAL1-CEN6-pGAL1-skHIS3 ADE2 NATr::16 mutated Rap1 sites 6bp gaps in chromosome 6 at position 192140 CEN4::klLEU2 in chromosome 6 at position 245710* | S4 |
| YAD86 | *MATa bar1-Δ KANr-pGAL1-CEN6-pGAL1-skHIS3 ADE2 NATr::16 Rap1 sites 35bp gaps 6bp gaps in chromosome 6 at position 192140 CEN4::klLEU2 in chromosome 6 at position 245710* | S4 |
| YAD97 | *MATa bar1-Δ KANr-pGAL1-CEN6-pGAL1-skHIS3 ADE2 NATr::16 Rap1 sites 6bp gaps in chromosome 6 at position 192140 CEN4::klLEU2 in chromosome 6 at position 245710* | S4 |

| | | |
|---|---|---|
| YAD128 | *MATa bar1-Δ KANr-pGAL1-CEN6-pGAL1-skHIS3 ADE2 NATr::16 mutated Rap1 sites 10bp gaps in chromosome 6 at position 192140 CEN4::klLEU2 in chromosome 6 at position 245710* | S4 |
| YAD130 | *MATa bar1-Δ KANr-pGAL1-CEN6-pGAL1-skHIS3 ADE2 NATr::16 mutated Rap1 sites 15bp gaps in chromosome 6 at position 192140 CEN4::klLEU2 in chromosome 6 at position 245710* | S4 |
| YAD138 | *MATa bar1-Δ KANr-pGAL1-CEN6-pGAL1-skHIS3 ADE2 NATr::16 mutated Rap1 sites 35bpgaps in chromosome 6 at position 192140 ura3-1::lacI-mCherry*::URA3 CEN4::klLEU2 in chromosome 6 at position 245710* | S4 |
| YAD139 | *MATa bar1-Δ KANr-pGAL1-CEN6-pGAL1-skHIS3 ADE2 NATr::16 mutated Rap1 sites 35bpgaps in chromosome 6 at position 192140 ura3-1::lacI-mCherry::URA3 CEN4::klLEU2 in chromosome 6 at position 245710* | S4 |
| YAD148 | *MATa bar1-Δ KANr-pGAL1-CEN6-pGAL1-skHIS3 ADE2 NATr::16 mutated Rap1 sites 20bp gaps in chromosome 6 at position 192140 CEN4::klLEU2 in chromosome 6 at position 245710* | S4 |
| YAD184 | *MATa bar1-Δ KANr-pGAL1-CEN6-pGAL1-skHIS3 ADE2 NATr::16 mutated Rap1 sites 30bp gaps in chromosome 6 at position 192140 CEN4::klLEU2 in chromosome 6 at position 245710* | S4 |
| YAD267 | *MATa hml::ADE1 hmr::ADE1 ade3::GALHO MATa::TetO-LEU2 0.5kb-CWH43::lacOpFX-TRP1 ura3::OsTIR1-URA3-LacI-mCherry-KanMx leu2::TetR-GFP-LEU2 Myo1-GFP::HPHBr* | 4A |
| YAD 302 | *MATa MATa hml::ADE1 hmr::ADE1 ade3::GALHO MATa::TetO-LEU2 0.5kb-CWH43::lacOpFX-TRP1 ura3::OsTIR1-URA3-LacI-mCherry-KanMx leu2::TetR-GFP-LEU2 Myo1-GFP::HPHB 16Rap1-6bp-gaps -at-chr.3-at-1670517::NATr* | 4A |
| YAD317 | *MATa MATa hml::ADE1 hmr::ADE1 ade3::GALHO MATa::TetO-LEU2 0.5kb-CWH43::lacOpFX-TRP1 ura3::OsTIR1-URA3-LacI-mCherry-KanMx leu2::TetR-GFP-LEU2 Myo1-GFP::HPHB no_Rap1-at-chr.3-at-1670517::NATr* | 4A |
| YAD319 | *MATa MATa hml::ADE1 hmr::ADE1 ade3::GALHO MATa::TetO-LEU2 0.5kb-CWH43::lacOpFX-TRP1 ura3::OsTIR1-URA3-LacI-mCherry-KanMx leu2::TetR-GFP-LEU2 Myo1-GFP::HPHB 16mutated-6bp-gaps-at-chr.3-at-1670517::NATr* | 4A |
| YAD319 | *MATa MATa hml::ADE1 hmr::ADE1 ade3::GALHO MATa::TetO-LEU2 0.5kb-CWH43::lacOpFX-TRP1 ura3::OsTIR1-URA3-LacI-mCherry-KanMx leu2::TetR-GFP-LEU2 Myo1-GFP::HPHB 16Rap1-30bp-gaps-at-chr.3-at-1670517::NATr* | 4A |
| Lev915 | *MATa MATa hml::ADE1 hmr::ADE1 ade3::GALHO MATa::TetO-LEU2 0.5kb-CWH43::lacOpFX-TRP1 ura3::OsTIR1-URA3-LacI-mCherry-KanMx leu2::TetR-GFP-LEU2 Myo1-GFP::HPHB smc2-AID-9myc::NATr* | 4A |

**Table S2: Rap1 array sequences with their contour lengths in bp and nm.** Rap1 binding sites are indicated in red. All arrays were initially created in a pUC19-BBB cloning vector where BamHI and BglII cloning sites (underlined) replace the pUC19 original polylinker: ---AGTGAATTGGGAC <u>GGATCC</u>TGATCA<u>AGATCT</u>AGCTTGGCGTAATCATGGT---.

| Construct | Length [bp] | Length [nm] | Sequence |
|---|---|---|---|
| 16Rap1-6bpGap | 274 | 93 | GGATCCGGTGTCTGGGTGTAAGGTGTATGGGTGTAGGATCTGGTGTCT<br>GGGTGTAAGGTGTATGGGTGTAGGATCTGGTGTCTGGGTGTAAGGTGT<br>ATGGGTGTAGGATCTGGTGTCTGGGTGTAAGGTGTATGGGTGTAGGAT<br>CTGGTGTCTGGGTGTAAGGTGTATGGGTGTAGGATCTGGTGTCTGGGT<br>GTAAGGTGTATGGGTGTAGGATCTGGTGTCTGGGTGTAAGGTGTATGG<br>GTGTAGGATCTGGTGTCTGGGTGTAAGGTGTATGGGTGTAAGATCT |
| 16Rap1-10bpGap | 302 | 103 | GGATCCGGTGTCTGGGTGTAAGGTGTATGGGTGTAGGATCTGAATGGT<br>GTCTGGGTGTAAGGTGTATGGGTGTAGGATCTGAATGGTGTCTGGGTG<br>TAAGGTGTATGGGTGTAGGATCTGAATGGTGTCTGGGTGTAAGGTGTA<br>TGGGTGTAGGATCTGAATGGTGTCTGGGTGTAAGGTGTATGGGTGTAG<br>GATCTGAATGGTGTCTGGGTGTAAGGTGTATGGGTGTAGGATCTGAAT<br>GGTGTCTGGGTGTAAGGTGTATGGGTGTAGGATCTGAATGGTGTCTGG<br>GTGTAAGGTGTATGGGTGTAAGATCT |
| 16Rap1-15bpGap | 337 | 115 | GGATCCGGTGTCTGGGTGTAAGGTGTATGGGTGTAGGATCTGATCCAT<br>ATGGTGTCTGGGTGTAAGGTGTATGGGTGTAGGATCTGATCCATATGG<br>TGTCTGGGTGTAAGGTGTATGGGTGTAGGATCTGATCCATATGGTGTC<br>TGGGTGTAAGGTGTATGGGTGTAGGATCTGATCCATATGGTGTCTGGG<br>TGTAAGGTGTATGGGTGTAGGATCTGATCCATATGGTGTCTGGGTGTA<br>AGGTGTATGGGTGTAGGATCTGATCCATATGGTGTCTGGGTGTAAGGT<br>GTATGGGTGTAGGATCTGATCCATATGGTGTCTGGGTGTAAGGTGTAT<br>GGGTGTAAGATCT |
| 16Rap1-20bpGap | 372 | 126 | GGATCCGGTGTCTGGGTGTAAGGTGTATGGGTGTAGGATCTGAACAAT<br>TCCATATGGTGTCTGGGTGTAAGGTGTATGGGTGTAGGATCTGAACAA<br>TTCCATATGGTGTCTGGGTGTAAGGTGTATGGGTGTAGGATCTGAACA<br>ATTCCATATGGTGTCTGGGTGTAAGGTGTATGGGTGTAGGATCTGAAC<br>AATTCCATATGGTGTCTGGGTGTAAGGTGTATGGGTGTAGGATCTGAA<br>CAATTCCATATGGTGTCTGGGTGTAAGGTGTATGGGTGTAGGATCTGA<br>ACAATTCCATATGGTGTCTGGGTGTAAGGTGTATGGGTGTAGGATCTG<br>AACAATTCCATATGGTGTCTGGGTGTAAGGTGTATGGGTGTAAGATCT |
| 16Rap1-25bpGap | 407 | 138 | GGATCCGGTGTCTGGGTGTAAGGTGTATGGGTGTAGGATCTGACGCTC<br>ACAATTCCATATGGTGTCTGGGTGTAAGGTGTATGGGTGTAGGATCTG<br>ACGCTCACAATTCCATATGGTGTCTGGGTGTAAGGTGTATGGGTGTAG<br>GATCTGACGCTCACAATTCCATATGGTGTCTGGGTGTAAGGTGTATGG<br>GTGTAGGATCTGACGCTCACAATTCCATATGGTGTCTGGGTGTAAGGT<br>GTATGGGTGTAGGATCTGACGCTCACAATTCCATATGGTGTCTGGGTGT<br>AAGGTGTATGGGTGTAGGATCTGACGCTCACAATTCCATATGGTGTCT<br>GGGTGTAAGGTGTATGGGTGTAGGATCTGACGCTCACAATTCCATATG<br>GTGTCTGGGTGTAAGGTGTATGGGTGTAAGATCT |
| 16Rap1-30bpGap | 442 | 150 | GGATCCGGTGTCTGGGTGTAAGGTGTATGGGTGTAGGATCTGATTATC<br>CGCTCACAATTCCATATGGTGTCTGGGTGTAAGGTGTATGGGTGTAGG<br>ATCTGATTATCCGCTCACAATTCCATATGGTGTCTGGGTGTAAGGTGTA<br>TGGGTGTAGGATCTGATTATCCGCTCACAATTCCATATGGTGTCTGGGT<br>GTAAGGTGTATGGGTGTAGGATCTGATTATCCGCTCACAATTCCATAT |

| | | | |
|---|---|---|---|
| | | | GGTGTCTGGGTGTAAGGTGTATGGGTGTAGGATCTGATTATCCGCTCACAATTCCATATGGTGTCTGGGTGTAAGGTGTATGGGTGTAGGATCTGATTATCCGCTCACAATTCCATATGGTGTCTGGGTGTAAGGTGTATGGGTGTAAGATCT |
| 16Rap1-35bpGap | 477 | 162 | GGATCCGGTGTCTGGGTGTAAGGTGTATGGGTGTAGGATCTGAAATTGTTATCCGCTCACAATTCCATATGGTGTCTGGGTGTAAGGTGTATGGGTGTAGGATCTGAAATTGTTATCCGCTCACAATTCCATATGGTGTCTGGGTGTAAGGTGTATGGGTGTAGGATCTGAAATTGTTATCCGCTCACAATTCCATATGGTGTCTGGGTGTAAGGTGTATGGGTGTAGGATCTGAAATTGTTATCCGCTCACAATTCCATATGGTGTCTGGGTGTAAGGTGTATGGGTGTAGGATCTGAAATTGTTATCCGCTCACAATTCCATATGGTGTCTGGGTGTAAGGTGTATGGGTGTAGGATCTGAAATTGTTATCCGCTCACAATTCCATATGGTGTCTGGGTGTAAGGTGTATGGGTGTAGGATCTGAAATTGTTATCCGCTCACAATTCCATATGGTGTCTGGGTGTAAGGTGTATGGGTGTAAGATCT |
| 2Rap1 | 29 | 10 | GGATCCGGTGTCTGGGTGTAAGGTGTATGGGTGTAAGATCT |
| 6Rap1-6bpGap | 99 | 34 | GGATCTGGTGTCTGGGTGTAAGGTGTATGGGTGTAGGATCTGGTGTCTGGGTGTAAGGTGTATGGGTGTAGGATCTGGTGTCTGGGTGTAAGGTGTATGGGTGTAAGATCT |
| 8Rap1-6bpGap | 134 | 46 | GGATCCGGTGTCTGGGTGTAAGGTGTATGGGTGTAGGATCTGGTGTCTGGGTGTAAGGTGTATGGGTGTAGGATCTGGTGTCTGGGTGTAAGGTGTATGGGTGTAGGATCTGGTGTCTGGGTGTAAGGTGTATGGGTGTAAGATCT |
| 12Rap1-6bpGap | 204 | 69 | GGATCCGGTGTCTGGGTGTAAGGTGTATGGGTGTAGGATCTGGTGTCTGGGTGTAAGGTGTATGGGTGTAGGATCTGGTGTCTGGGTGTAAGGTGTATGGGTGTAGGATCTGGTGTCTGGGTGTAAGGTGTATGGGTGTAGGATCTGGTGTCTGGGTGTAAGGTGTATGGGTGTAGGATCTGGTGTCTGGGTGTAAGGTGTATGGGTGTAAGATCT |

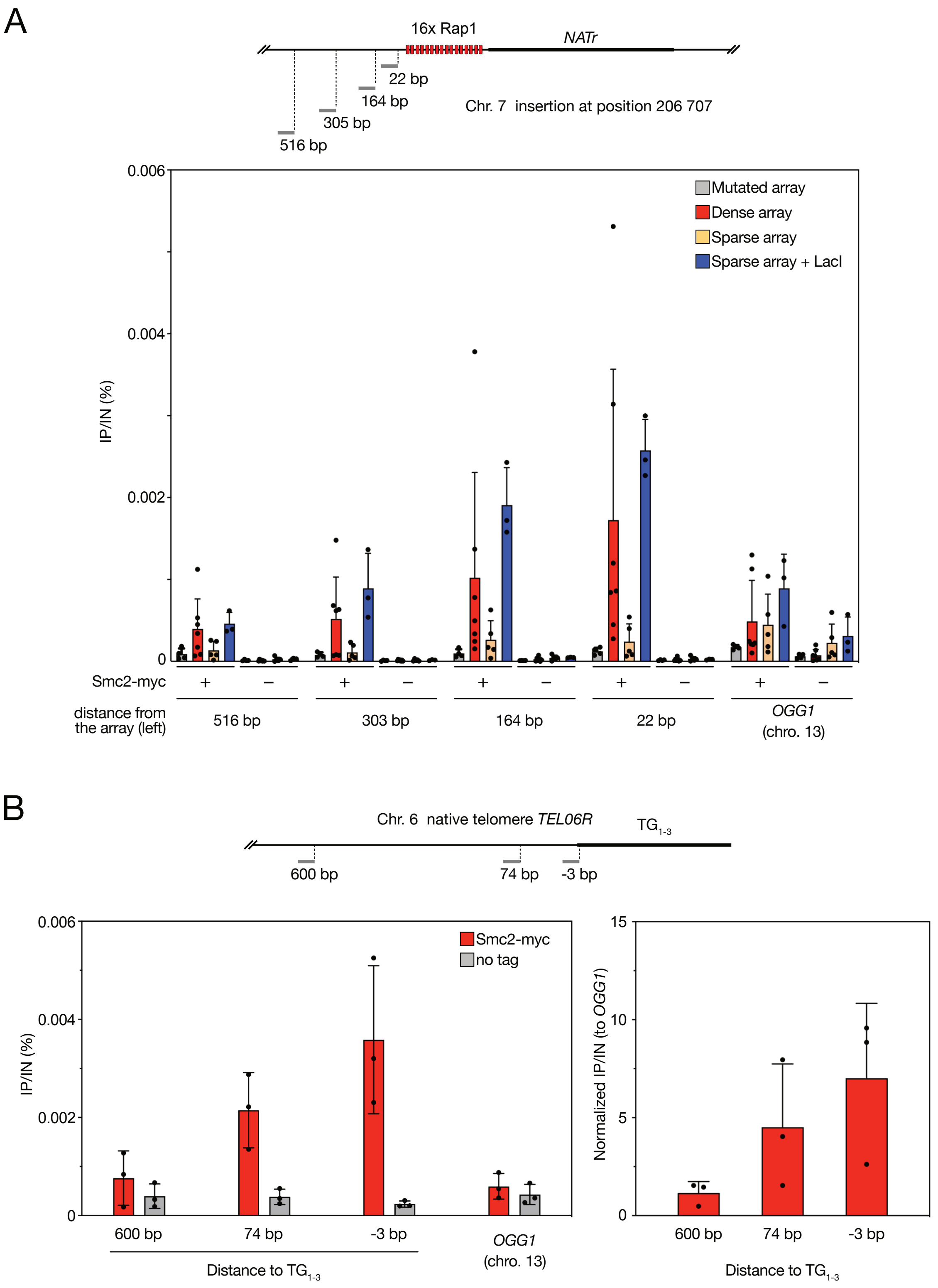


**Fig. S1. ChIP analysis of Smc2-Myc in cells synchronized in late anaphase. (A)** Individual IP/Input values from which the relative enrichment at Rap1 arrays shown in Fig.1C is derived**.** Error bars indicate standard deviation **(B)** Condensin enrichment at native telomere *TEL06R.* Left panel: Individual IP/Input values. Right panel: enrichment relative to an ectopic internal position. The bars represent mean IP/Input values; error bars indicate standard deviation over 3 biological replicates.

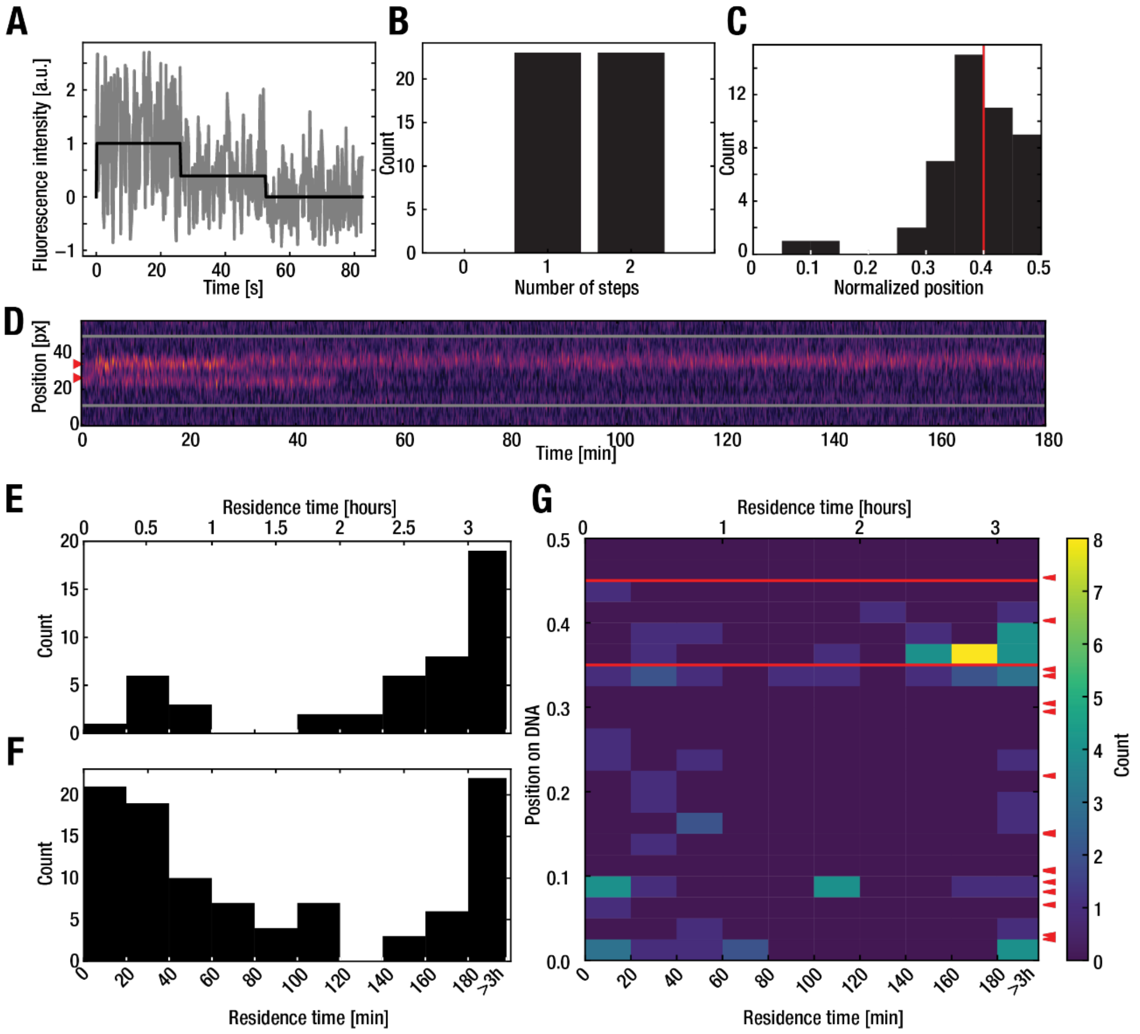


**Fig. S2. Validation of binding efficiency, specificity, and residence time. (A-C)** Bleaching assay under Rap1 DNA-binding conditions (see methods). **(A)** Representative bleaching curve of two Rap1-JF646 proteins bound to DNA (see methods). The raw fluorescence intensity is shown in grey and the steps are shown in black. **(B)** Histogram of identified bleaching steps (N=46). **(C)** Histogram of the positions of bleached Rap1-JF646 in (B). Positions are normalized by the end-to-end length. Distances are shown from 0-0.5 because we cannot distinguish directionality of the DNA molecules. **(D-G)** Residence time assay under imaging conditions (see methods). **(D)** Representative example of a kymograph that shows a Rap1 protein with a long and short residence time. The red arrows indicate the position of the Rap1 binding site located at 0.4 distance along the DNA (because we cannot distinguish top from bottom and our binding site is located asymmetrically along the DNA, we indicate both 0.4 and the mirrored 0.6 position, which is coincidentally located near several potential Rap1 sites within the i95-cosmid sequence). **(E)** Histogram of residence time measured for Rap1 proteins near the binding site position. Last bin (>3h) represents all proteins that were still present at the end of acquisition time. **(F)** Histogram of residence time measured for all off-target bound Rap1 proteins. **(G)** 2D histogram of the residence time (x-axis) and binding position (y-axis). Red lines indicate the positional bins from which on-target binding was aggregated to generate the histogram in (D). Red arrows indicate sites partially matching Rap1 consensus site (5'-RRKGNKYGGRTKY-3') within the i95-cosmid sequence.

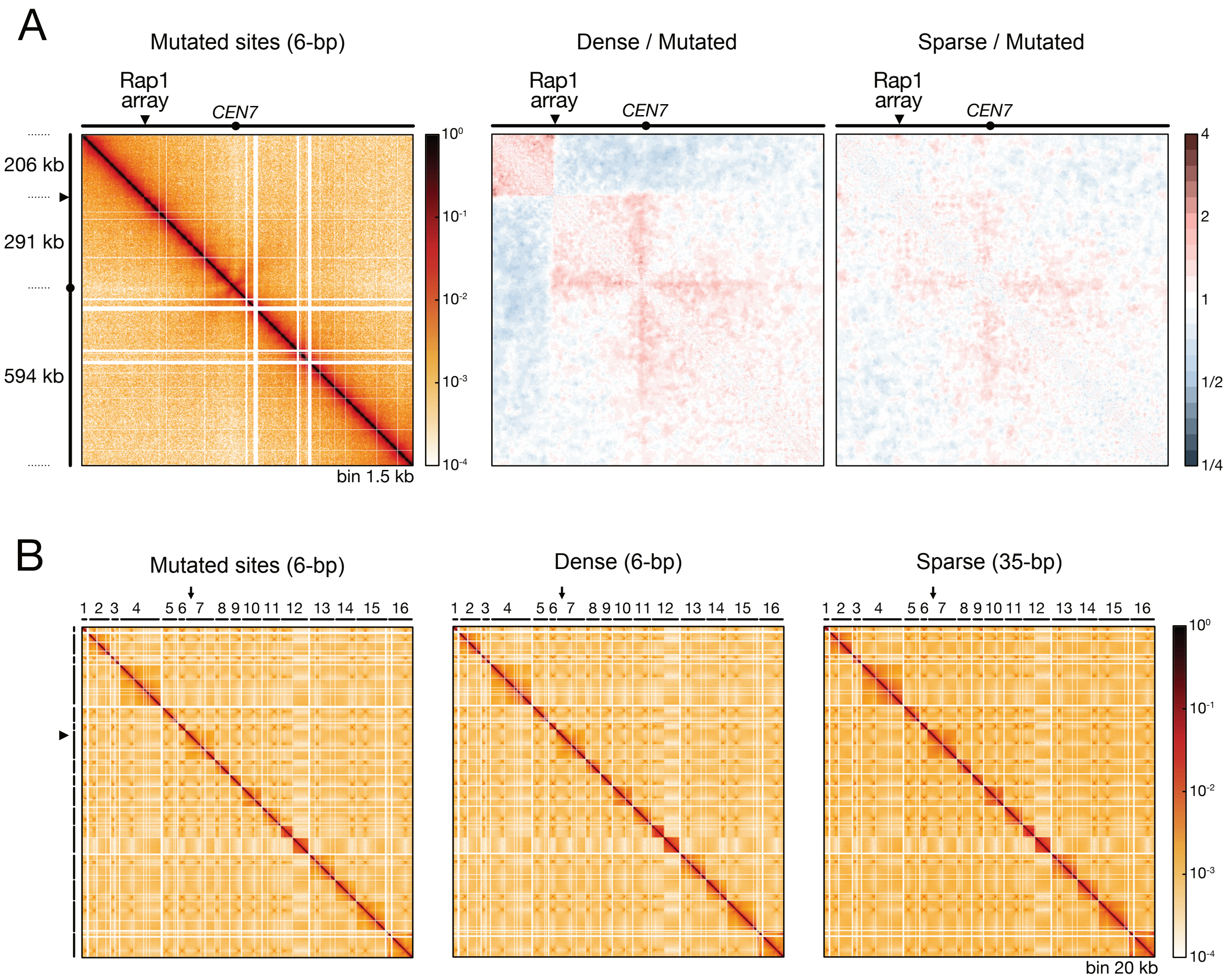


**Fig. S3. MicroC at Rap1 arrays.** (A) MicroC contact map of chromosome 7 where a control array of 16 mutated sites is inserted at position 206 707. Cells are synchronized in late anaphase. The ratios were determined using the Serpentine tool. We note that dense and sparse Rap1 arrays make frequent contact with the proximal native chromosome end. Unlikely to be related to condensin activity, these *cis*-contacts may stem instead from Rap1-dependent direct protein-protein interactions known to cluster telomeres[1]. (B) Whole-genome contact maps.

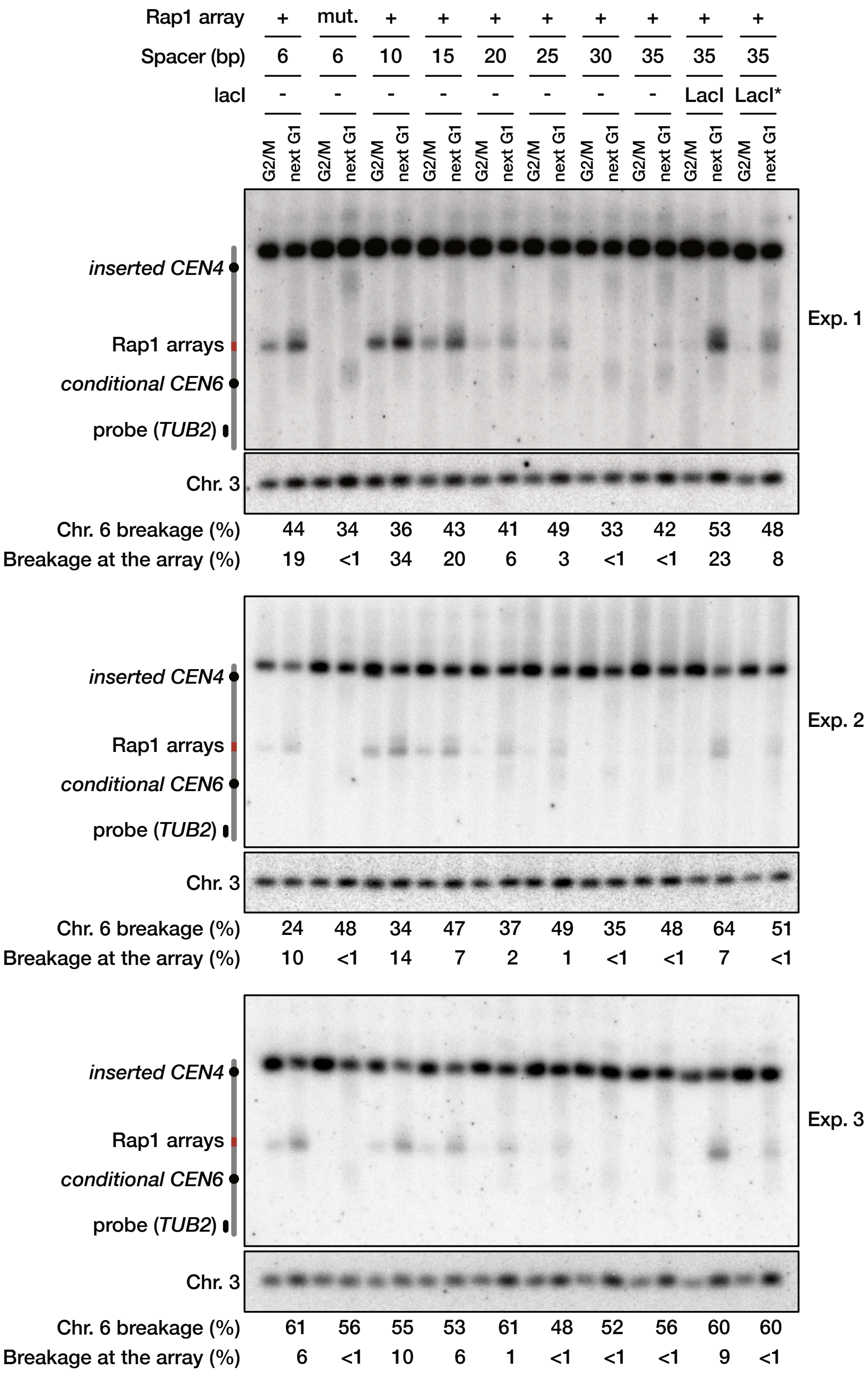


**Fig.S4. Dicentric chromosome breakage.** Individual gels from which the preferential dicentric breakage at Rap1 arrays shown in Fig.4C is derived.

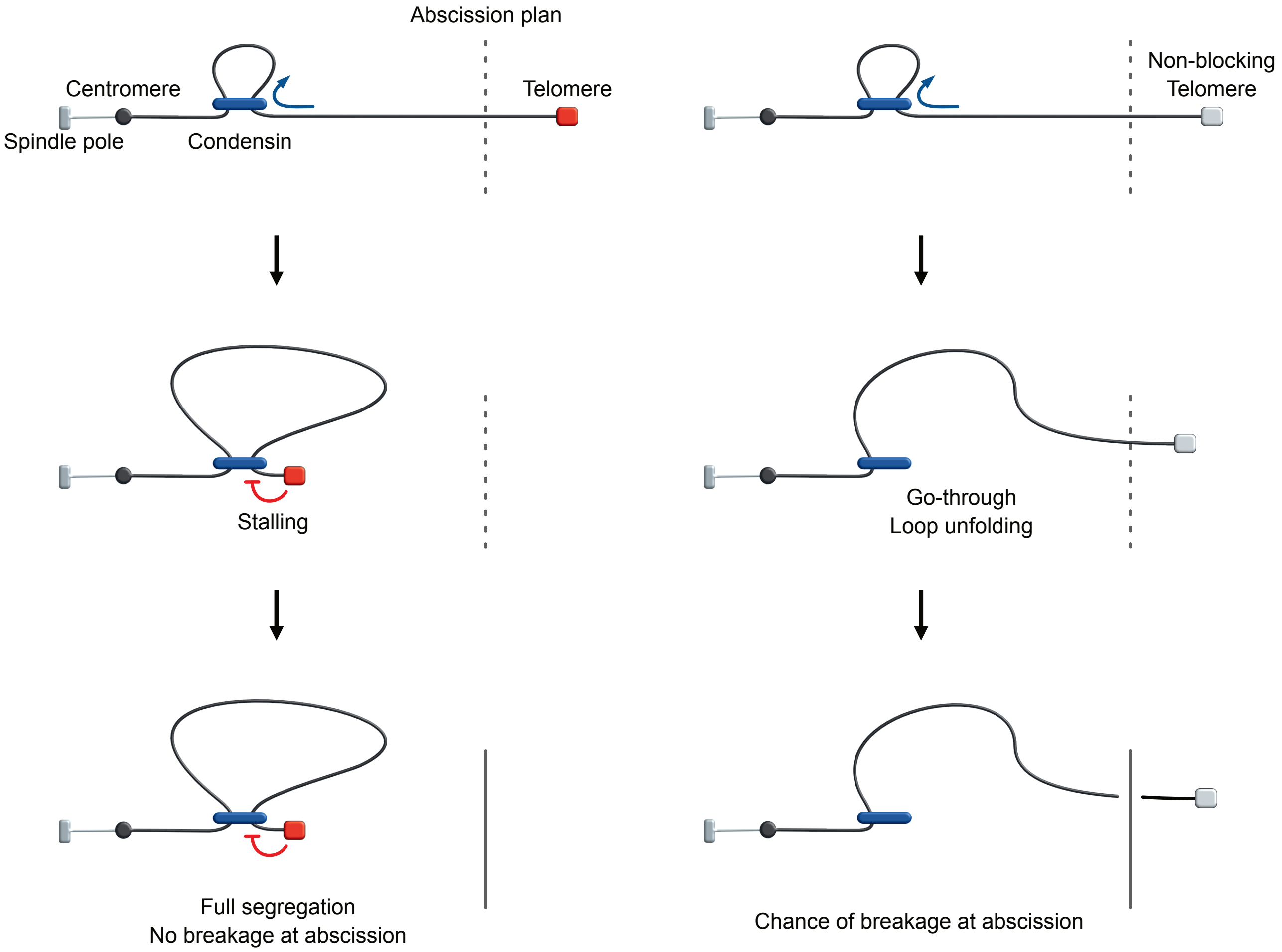


**Fig.S5. Working model for a role of condensin stalling at native telomeres.**
